# Multiplex qPCR assays for detection of two imperiled anuran species, *Anaxyrus californicus* and *Spea hammondii*, from environmental DNA

**DOI:** 10.1101/2022.12.20.521282

**Authors:** Torrey W. Rodgers, Robert E. Lovich, James A. Walton, Daniel J. Prince, Bernardo R. Gonzalez, H. Bradley Shaffer, Karen E. Mock

## Abstract

We developed a species-specific, quantitative PCR assay multiplex for the detection of two threatened and endangered anuran species, arroyo toad (*Anaxyrus californicus)* and western spadefoot (*Spea hammondii*), from environmental DNA (eDNA). Both species have experienced range wide declines over the last century, mostly as a result of habitat loss. Given these declines and their cryptic life histories, improved tools for detecting and monitoring both species are needed. Species-specificity and sensitivity of assays were empirically tested in the lab, and the multiplexed assays were validated with field-collected eDNA samples. Both assays were species-specific, sensitive, and effectively detected the target species from streams, rivers and ponds, although detection was imperfect. These assays provide an efficient and economical tool that will aid in the detection, monitoring, management, and conservation of these at-risk species.

## INTRODUCTION

The arroyo toad (*Anaxyrus californicus*) and western spadefoot (*Spea hammondii*) are native to the state of California in the United States, and the adjacent Mexican state of Baja California. Both species have experienced dramatic population declines in the past century, primarily due to loss of breeding habitat (Thomson et al. 2016). Because both species use ephemeral or intermittent water bodies for breeding, they are susceptible to demographic declines and inbreeding depression due to climate change and drought (Neal et al. 2020). Arroyo toads are breeding habitat specialists in slow-flowing, low-gradient, sand or gravel-bottomed streams (Thomson et al., 2016). They have been extirpated from as much as 75% of their historical range in California (Jennings and Hayes 1994, USFWS 2014) and were listed as endangered under the US Endangered Species Act in 1994 (Federal Register 1994). Western spadefoot are more generalized in their breeding requirements, and will utilize natural vernal pools and constructed cattle and other ponds that are predator free and hold water for at least 36 days (Thomson et al. 2016). This species has also seen dramatic population declines and is estimated to be absent from greater than 30% of its historic habitat in northern and central California, and as much as 80% in southern California (Jennings and Hayes 1994, Fisher and Shaffer 1996, Thomson et al. 2016). Western spadefoot are currently listed as a Priority I Species of Special Concern in California (Thomson et al. 2016), and a formal review for listing under the Endangered Species Act has been initiated by the U.S. Fish and Wildlife Service (Neal et al. 2018, 2020).

Efficient monitoring of species’ distributions is essential to inform management and conservation efforts and doing so for cryptic species like arroyo toad and western spadefoot can be challenging. To this end, we designed multiplexed, species-specific, quantitative PCR (qPCR) assays for the detection of both species from environmental DNA (eDNA) samples. eDNA sampling is a molecular technique for detecting trace amounts of DNA released by organisms into the water column. eDNA techniques are highly sensitive and can be more efficient and economical than traditional sampling (Fediajevaite et al. 2021). The potential to use eDNA for detection of arroyo toad and western spadefoot will allow for more efficient monitoring of these imperiled species.

## METHODS

### Assay Design

For assay design, we sequenced mitochondrial DNA from tissue samples from across the ranges of both species (Fig 1). For arroyo toad, we sequenced 627 base pairs (bp) of the mitochondrial cytochrome b (cytb) gene from 74 tissue samples, including 60 from California, USA and 14 from Baja California, Mexico (Table S1, Fig 1a). For western spadefoot, we sequenced 607 bp of cytb, as well as 530 bp of the mitochondrial 16S rRNA gene (16S) from 42 tissue samples from across California (Table S2, Fig 1b). We generated sequence data from two mitochondrial genes for the western spadefoot because highly distinct, reciprocally monophyletic northern and southern clades occur within *S. hammondii* (Neal et al. 2018). Given this relatively deep differentiation, we thought it prudent to explore both a more conserved (16S) and a more variable (cytb) gene region for designing an assay capable of detecting both *S. hammondii* clades, in case the cytb gene region proved too variable. We also sequenced 43 tissue samples from 20 potentially sympatric amphibian and turtle species for cytb or 16S (Table S3) and obtained additional non-target species sequences from GenBank (Table S4) to design species-specific assays with a low probability of amplifying DNA from sympatric species.

**Figure 1.**
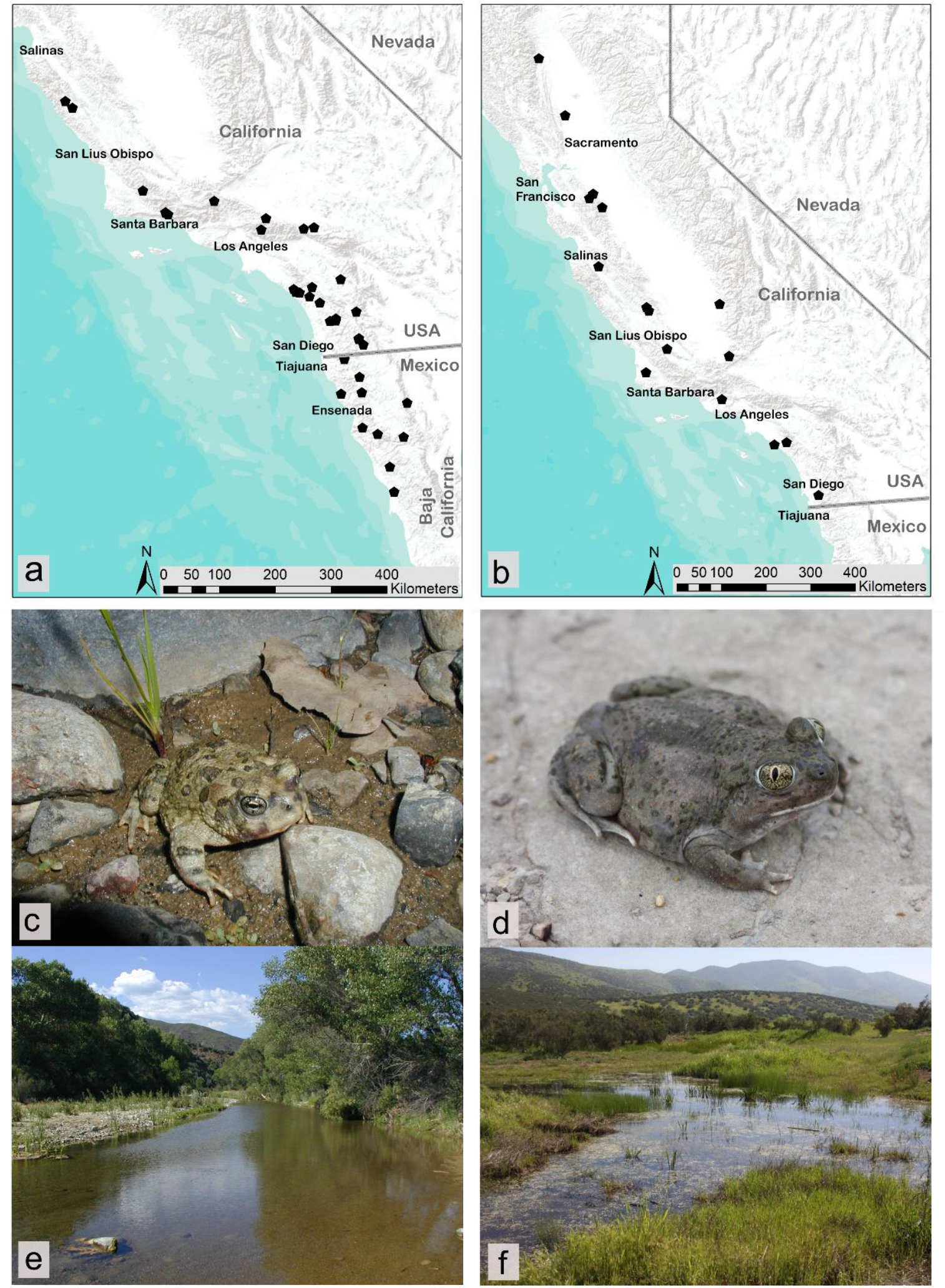
a) Geographic locations of target-species tissue samples sequenced and used for design of species-specific qPCR assays for *Anaxyrus californicus*, and b) Geographic locations of target-species tissue samples sequenced and used for design of species-specific qPCR assays for *Spea hammondii*. c) Adult *Anaxyrus californicus*. d) Adult *Spea hammondii*. e) *Anaxyrus californicus* habitat, Los Padres National Forest, California. f) *Spea hammondii* habitat, San Diego County, California.

Sequence data from all species were aligned using Sequencher software v5.2 (Gene Codes, Ann Arbor, MI), and species-specific primers were generated with the online tool DECIPHER (Wright et al. 2014). For Arroyo toad, we initially designed and tested three candidate primer sets targeting cytb (Table S5). For western spadefoot, we initially designed and tested three candidate primer sets, two targeting cytb, and one targeting 16S (Table S5). Candidate primers were initially tested with SYBR^®^ green qPCR, and one primer set for each species was chosen for further testing based on specificity and efficiency, as described below. We then used ABI primer express software v3.0.1 (Applied Biosystems, Foster City, CA) to design TaqMan™ Minor-Groove-Binder (MGB) fluorescent qPCR probes for each primer set with as many bp mismatches with non-target species as possible.

### Specificity testing

Specificity testing included both *in silico* and *in vitro* components. To evaluate primer specificity *in silico*, we used NCBI Primer-BLAST (Ye et al. 2012) against the NCBI refseq non-redundant (nr) database to assess the likelihood that species occurring within or near the ranges of both species could potentially cross-amplify and produce false positives. Primer-BLAST stringency settings required the primers have at least 2 total mismatches to unintended targets, including at least 1 mismatch within the last 5 bps of the 3’ end, and ignored targets that had 6 or more mismatches to the primer.

In the laboratory, we initially evaluated the specificity of our six candidate primer sets using SYBR^®^ green qPCR to find a primer set with optimal specificity for each target species. Each primer set was tested with 10 tissue DNA extracts from different individuals of each of the target species, as well as DNA extracts from at least two individuals from each sympatric non-target anuran species (n=9 species; Table S3). qPCR reactions included 7.5 μl Power SYBR^®^-Green Mastermix (Thermo-Fisher Scientific), 900 nM of each primer, and 0.1 ng of template DNA in a total reaction volume of 15 μl. Cycling conditions were 95° C for 10 minutes followed by 45 cycles of 95° C for 15 seconds and 60° C for one minute, followed by a melt curve to check for primer-dimer formation or off-target amplification. After one optimal primer set was chosen for each species (primer sets ANCA3 and SPHA2; see results), we then tested all four primers in multiplex with SYBR^®^-Green qPCR on tissue samples of both target species with a melt curve to check for potential primer-dimer formation in the multiplex.

Primer sets and MGB probes were then tested in TaqMan™ qPCR. We tested DNA extracts from ten individuals from each target species, two DNA extracts from each non-target anuran species (n=9 species), and one or two DNA extracts from each non-target urodele (salamander or newt) (n= 8 species) and turtle species (n=4 species; Table S3). qPCR reactions included 7.5 μl TaqMan™ Environmental Mastermix 2.0 (Thermo-Fisher Scientific), 900 nM of each primer, 250 nM of probe, and 0.1 ng of template DNA in a total reaction volume of 15 μl. Cycling conditions were 95° C for 10 minutes followed by 45 cycles of 95° C for 15 seconds and 60° C for one minute.

### Primer concentration optimization

Primer concentrations were optimized by running four concentrations of each forward and reverse primer (100 nM, 300 nM, 600 nM, and 900 nM), each in triplicate, with 0.004 ng of target-species tissue DNA per reaction. Primer concentrations with the greatest peak fluorescence and lowest C_t_ value were used for subsequent analyses (Table 1).

**Table 1.**
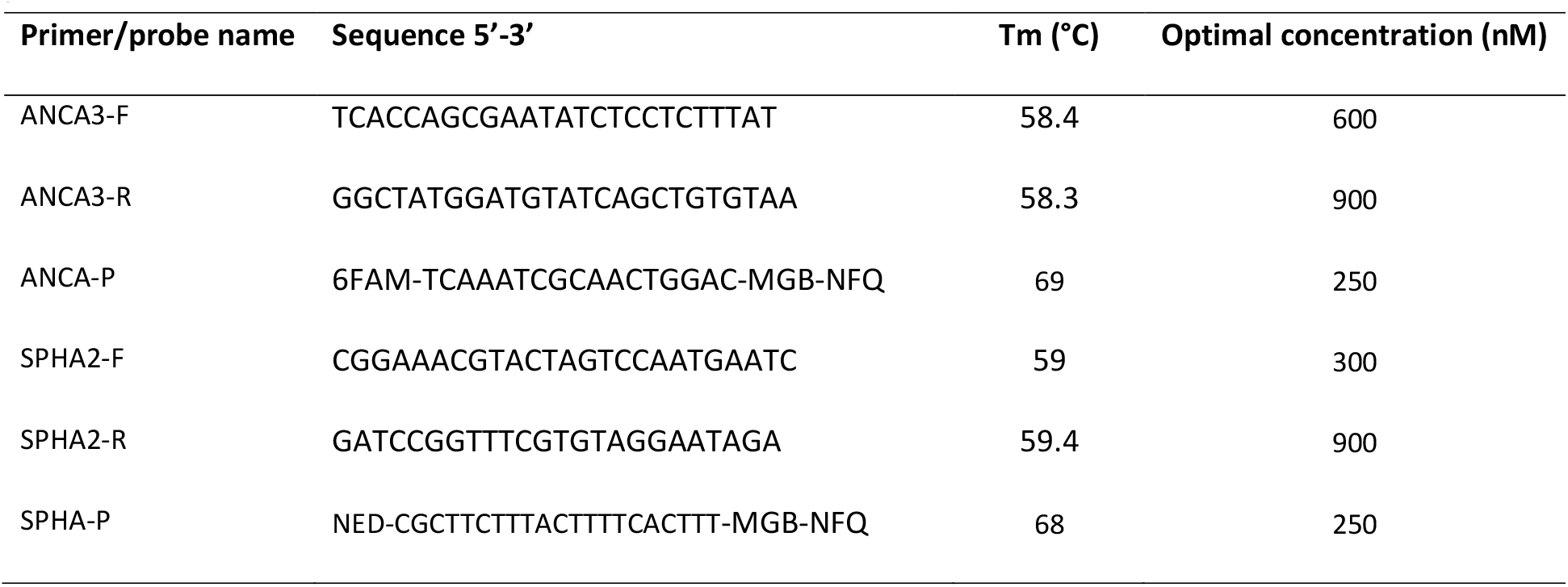
Primers and probes used in the multiplex assay for detection of *Anaxyrus californicus* and *Spea hammondii* from environmental DNA. F and R refer to forward and reverse primers; P refers to the MGB probe.

### Sensitivity testing

To test assay sensitivity, we used MiniGene™ synthetic plasmids containing the assay sequences for each species. Plasmids were suspended in 100 μL of IDTE buffer (10 mM Tris, 0.1 mM EDTA), linearized by digestion with the enzyme Pvu1, and then purified with a PureLink™ PCR Micro Kit (Invitrogen, Carlsbad, CA) following standard protocols. The resulting products were quantified on an Invitrogen™ Qubit^®^ fluorometer, and estimated concentrations were converted to copy number based on molecular weight (Wilcox et al. 2013). These products were then diluted in IDTE buffer to create quantities of 2, 5, 10, 20, 50, and 100 copies/reaction of each plasmid, and each quantity was analyzed in ten qPCR replicates with the multiplex assay to determine assay sensitivity.

### Field validation

We field tested our assays using eDNA samples collected from water bodies where our target species had been documented within the prior four years (Hitchcock et al. 2022). We did not conduct exhaustive traditional surveys for our target species at the time of eDNA sampling at all sites. However, brief visual observations were made by eDNA technicians during sampling, and at a subset of sites, traditional visual, dipnet, or seine surveys were conducted by permitted individuals with expertise in morphological identification of our target species either during or shortly before eDNA sampling.

For arroyo toad, eDNA samples were collected from Marine Corps Base Camp Pendleton (MCBCP) in 2019 (n=8 total samples from seven sites) and 2022 (n=24 samples from eight sites), and from the Los Padres National Forest (LPNF) in Ventura County (n=15 samples from five sites) and Santa Barbara County (n=18 samples from 6 sites) in 2021. In 2019, we collected a single eDNA sample from each of seven sites, and duplicate eDNA samples from one site. In 2021 and 2022 triplicate eDNA samples were collected from each site (Table S6). For western spadefoot, we collected samples from U.S. Army Base Fort Hunter Liggett (FHL; n=23 samples from eight sites) in 2019, and from MCBCP (n= 12 total samples from four sites), U.S. Marine Corps Air Station Miramar (MCASM; n=3 samples from one site), and 2 private ranches in Santa Barbara County (n=15 samples from five sites) in 2021. All western spadefoot eDNA samples were collected in triplicate from each site, except for one where duplicate samples were collected (Table S7).

Environmental DNA samples were collected following the protocol outlined in Carim et al. (2016). Up to five liters of water were pumped through a 1.5 μm glass microfiber filter using a peristaltic pump in the field, and filters were stored in silica desiccant at -70 °C until laboratory processing. We additionally collected and analyzed ‘field blank’ filter samples (n=8) consisting of one liter of distilled water brought from the lab and filtered in the field with the same sampling equipment to act as a negative control for potential field-based between-sample contamination. Samples were extracted in a dedicated room using the DNeasy Blood & Tissue Kit (Qiagen, Inc. Valencia, CA, USA) with a modified protocol described in the supplementary material (Appendix I). Each round of eDNA extraction (n=10) included one ‘extraction blank’ negative control consisting of a clean, unused filter. All samples were analyzed in triplicate qPCR with 4 μl of eDNA extract, 7.5 μl TaqMan™ Environmental Mastermix 2.0 (Thermo-Fisher Scientific), optimized primer and probe concentrations (Table 2), and cycling conditions of 95° C for 10 minutes followed by 45 cycles of 95° C for 15 seconds and 60° C for one minute. All reactions also included a TaqMan™ Exogenous Internal Positive Control (Thermo-Fisher Scientific) to monitor for PCR inhibition, and each qPCR run included six ‘no template’ negative control reactions to monitor for cross-contamination. Samples were considered as “positive” when at least one of three qPCR replicates amplified prior to 45 PCR cycles. To further mitigate cross-contamination, all qPCR reactions were set up under a dedicated PCR hood sterilized with UV radiation prior to each qPCR run (Goldberg et al. 2016).

**Table 2.** Summary of environmental DNA sampling for arroyo toad *Anaxyrus californicus* and western spadefoot *Spea hammondii* for field validation of quantitative PCR assays.

| Year | Number of sites/samples | Number of sites with visual observation | Number of sites/samples with eDNA detection | % of sites/samples with eDNA detection at sites with visual observation |
| --- | --- | --- | --- | --- |
| **A. californicus** |  |  |  |  |
| 2019 | 7 / 8 | 6 | 6 / 7 | 100% / 100% |
| 2021 | 11 / 33 | 0 | 0 / 0 | Na / Na |
| 2022 | 8 / 24 | 8 | 5 / 11 | 63% / 46% |
| Total | 26 / 65 | 14 | 11 / 18 | 79% / 58% |
| **S. hammondi** |  |  |  |  |
| 2021 | 18 / 53 | 9 | 11 / 30 | 89% / 85% |

## RESULTS

### Assay design

The final species-specific assays we developed target a 120 bp and 147 bp fragment of cytb in arroyo toad and western spadefoot, respectively (Table 1). Ultimately, the more conserved 16S region was not necessary in designing an assay to detect both western spadefoot clades. Both cytb assays perfectly matched all available sequences from both target-species at both primers and the probe.

### Specificity testing

*In silico* evaluation of primer specificity with NCBI Primer-BLAST for arroyo toad identified no species with the potential to cross-amplify. For western spadefoot, Primer-BLAST identified several species with the potential to cross-amplify (Table S8). Of the taxa identified, just one occurs in North America: the congener *Spea bombifrons*, which does not occur in California or Baja California. All 32 *S. bombifrons* cytb sequences present on Genbank (accessed Nov 7, 2022), however, possess a terminal mismatch at the 3’ end of primer SPHA2-R such that cross-amplification is unlikely.

The *in vitro* evaluation of primer specificity with SYBR^®^ Green qPCR for arroyo toad determined of the three primer sets tested, ANCA3 was superior with regards to specificity. For primer set ANCA1 and ANCA2, tissue DNA from multiple non-target species amplified within 7-10 Cycles of arroyo toad tissue samples, a range which may not confer complete specificity once TaqMan™ probe is added. For primer set ANCA3 however, amplification was observed in just one non-target tissue sample from *Rana boylii*, and this sample amplified >19 cycles later than arroyo toad tissue samples, a range which is more than suitable for specificity once a TaqMan™ MGB probe is added. No other non-target sample amplified with primer set ANCA3, and all arroyo toad samples amplified with a mean C_t_ of 24.169. Additionally, the melt curve produced a single sharp peak indicating no primer-dimer formation or off-target amplification. Thus, we proceeded with primer set ANCA3 for further TaqMan™ qPCR testing.

For western spadefoot, *in vitro* evaluation of primer specificity with SYBR^®^ Green qPCR indicated that all three primer sets were species-specific amongst sympatric amphibian species. For all three primer sets, all non-target tissue samples either did not amplify, or amplified > 14 cycles later than target samples, a range which is more than suitable for specificity once a TaqMan™ MGB probe is added. We chose primer set SPHA2 to proceed with further TaqMan™ qPCR testing because SPHA2 had the lowest mean C_t_ for western spadefoot tissue samples (23.29) and displayed the highest peak fluorescence of the three primer sets tested. For SPHA2, all non-target samples amplified >16 cycles later than target samples or did not amplify at all, and the melt curve produced a single sharp peak indicating no primer-dimer formation or off-target amplification. In the SYBR^®^ Green test with both primer sets ANCA3 and SPHA2 in multiplex, the melt curve produced a single sharp peak indicating no primer-dimer formation between the 4 primers in the multiplex.

The *in vitro* evaluation of assay specificity with inclusion of TaqMan™ MGB probes indicated that both assays are species-specific amongst the sympatric amphibian samples tested, amplifying DNA from only the target species. For arroyo toad, all target samples amplified with a mean C_t_ of 27.42. For western spadefoot, all target samples amplified with a mean C_t_ of 28.07. No amplification of non-target tissue DNA was observed with either TaqMan™ assay.

### Sensitivity testing

For both species, 100% of qPCR replicates amplified down to a template concentration of 10 copies per reaction. For *A. californicus* 80% (8 of 10) of reactions containing 5 copies per reaction amplified, and 50% (5 of 10) of reactions containing 2 copies per reaction amplified. For *S. hammondii*, 90% (9 of 10) of reactions containing 5 copies per reaction amplified, and 60% (6 of 10) of reactions containing 2 copies per reaction amplified.

### Field validation

For arroyo toad, we collected 65 total eDNA samples at 26 sites from 10 rivers and streams (Table 2, Table S6) with recent documentation of species presence (Hitchcock et al. 2022). At 14 of the 26 sites, arroyo toad larvae and/or juveniles were observed either at the time of eDNA sampling (n=12) or by US Geological Survey (USGS) biologists conducting traditional surveys 2 weeks prior to our eDNA sampling (n=2; Robert Fisher, personal communication). All visual observations of larvae/juveniles were from MCBCP in 2019 and 2022. No arroyo toads of any life stage were visually observed during eDNA sampling in the LPNF during 2021, although the individual conducting eDNA sampling in 2021 was not an expert in the identification of arroyo toad larvae, which can be both cryptic and difficult to differentiate from other sympatric anuran larvae.

Our assay detected arroyo toad eDNA in 18 samples from 11 sites in 4 different rivers/streams (Table 2, Table S6). All eDNA detections were from sites where arroyo toad larvae or juveniles were visually observed at the time of eDNA sampling, or by USGS crews previous to eDNA sampling. In sampling at MCBCP in 2019, arroyo toad eDNA was detected in all samples where larvae or juveniles were observed. Arroyo toad eDNA was not detected in any eDNA water samples collected from the LPNF in 2021. For samples collected from MCBCP in 2022, eDNA was detected from 11/24 samples at 5/8 sites where larvae were observed at the time of sampling or by USGS biologists 2 weeks prior. eDNA failed to detect arroyo toad presence at 3/8 sites where they were visually confirmed as present (Table 2).

For western spadefoot, we collected 53 total eDNA samples at 18 sites from 13 ponds with historical documentation of species presence. At 9 of the 18 sites sampled, western spadefoot larvae were visually observed at the time of eDNA sampling. Sites with visual observation included three at FHL, two sites on private ranches in Santa Barbara County, and all sites at MCBCP (Table 2, Table S7). Our assay detected western spadefoot in 30 eDNA samples at 11 sites from seven lakes/ponds. This included 8 of the 9 sites where western spadefoot larvae were observed at the time of eDNA sampling. At one pond in Santa Barbara County, two western spadefoot larvae and one nearly metamorphosized individual were netted, but eDNA was not detected. Additionally, eDNA was detected at 3 sites from a single pond on FHL where western spadefoot larvae were observed in 2017 but were not observed at the time of eDNA sampling in 2019 (Table S7).

## DISCUSSION

We designed and validated species-specific qPCR assays for the detection of arroyo toad and western spadefoot, two at-risk anuran species of conservation concern, from eDNA. These assays are highly sensitive at detecting DNA from the target species and performed reasonably well in detecting both species from eDNA field samples at sites where they were observed at the time of or shortly before eDNA sampling. Although these species are not known to occur in the same aquatic habitats due to their unique breeding strategies, we designed the assays to be run in multiplex for those conducting surveys for both species.

We demonstrated that both of our assays are species-specific among all sympatric amphibian and turtle species by using reference sequences from sympatric species in the assay design, and empirically testing for cross-amplification against tissue DNA from sympatric species in the lab. However, we cannot say with certainty whether our assays might amplify DNA from non-sympatric, but closely related congeneric species. *Anaxyrus californicus* and *Anaxyrus mexicanus* were historically considered subspecies of *Anaxyrus microscaphus* but were later split into three species (Gergus 1998). At the time of writing, only one cytb reference sequence for *A. microscaphus* was available on Genbank (accession # L10978). This sequence only overlaps with the reverse primer of our assay but possesses a terminal 3’ mismatch with the primer, making cross-amplification unlikely. At the time of writing, no cytb reference sequences were available for *A. mexicanus*. The geographically nearest non-sympatric *Anaxyrus* species, *Anaxyrus punctatus*, has many cytb sequences available on Genbank, and all contain many mismatches with our primers, making cross-amplification unlikely. For our western spadefoot assay, many mismatches were observed with reference sequences of *Spea multiplicata*. Our assay had just one mismatch with all 32 Genbank reference sequences for *Spea bombifrons*. However, this was a terminal 3’ mismatch with the reverse primer, making cross-amplification unlikely. Our assay could potentially be modified at this single position to be used to detect *Spea bombifrons*. At the time of writing, no cytb reference sequences were available on Genbank for the congener *Spea intermontana*. Neal et al. (2018) found that the ND4 mitochondrial sequence of the southern clade of *S. hammondii* was more similar to that of *S. intermontana* than to the northern clade of *S. hammondii*, possibly due to a historical mitochondrial replacement event. Thus, it seems probable that our assay may amplify *S. intermontana* DNA, however additional sequence data from *S. intermontana* and laboratory testing would be needed to determine if this is the case.

Locating appropriate sites for field validation of our arroyo toad assay was difficult due to the severe drought in California in 2020-2022. In 2019, our sampling sites experienced above average rainfall, however we collected just a few validation samples in 2019 as our assay development was in the pilot phase. In 2021, we attempted to conduct more extensive validation sampling for arroyo toad on the LPNF in Ventura and Santa Barbara counties, as this region includes designated critical habitat for arroyo toad and well-documented breeding populations. Because precipitation in the winter of 2020-2021 was far below average, we began conducting our eDNA sampling in May, slightly prior to the peak breeding season in a ‘normal’ year for these higher elevation populations (Samuel Sweet, personal communication). However, even at that time some sites had low water, and no arroyo toad larvae were observed during our field sampling. By June, during the peak breeding season in a ‘normal’ year in these populations, most breeding sites were completely dry. We did not detect arroyo toad eDNA from any water samples from the LPNF in 2021. As the drought continued in the 2022 breeding season, we chose a single river to complete our field validation of the arroyo toad assay; the Santa Margarita River on MCBCP in San Diego County. We chose this river because it is one of the only arroyo toad breeding areas with perennial water flow and is known to be one of the only locations where arroyo toad breeding is often successful in dry years.

As eDNA sampling most commonly utilizes water for detection of aquatic species, drought conditions will lessen the utility of using eDNA to monitor aquatic breeding toads and ecologically similar species. We did, however, have one terrestrial arroyo toad detection from the LPNF in 2021, even though no breeding was observed, and all water eDNA samples were negative. This single positive detection came from a toad scat sample. We collected 11 putative toad scat samples from the LPNF in 2021 as a pilot study and ran our arroyo toad assay on them (data not shown). We detected arroyo toad eDNA from 1/11 scats. However, it is likely that many of the scats were from congeneric *A. boreas*, which is common in the study area and has scat that is morphologically indistinguishable from *A. californicus*. Although we only had a single observation, this suggests that our qPCR assays may have utility for monitoring adult toads using scat samples, especially in dry years when water for typical eDNA sampling is scarce and toads are not breeding. More work is needed to determine the utility of this approach.

For arroyo toad, we detected eDNA at most locations where larvae or juveniles were observed during, or shortly before (within 2 weeks) of eDNA sampling. In our 2019 sampling, we detected arroyo toads in every sample where larvae or juveniles were observed. This was not surprising, as 2019 was an excellent breeding year with above average precipitation, and larvae and juvenile toads were abundant at sampling sites. Conversely, 2022 was a very dry, poor breeding year, and only a very few arroyo toad larvae were observed in our sampling sites on the Santa Margarita River. In 2022 we detected arroyo toad eDNA from 5 of 8 sites where larvae were observed during or within 2 weeks prior to sampling. At one site, a water weir broke above the site hours before sampling, causing a sudden increase in stream flow, and likely diluting or washing away any eDNA. Thus, our failure to detect eDNA at that site, despite larvae having been observed 2 weeks previous, is understandable. However, at two other sites we also failed to detect arroyo toads with eDNA, even though larvae were observed at the time of sampling. Additionally, in 2022, arroyo toad eDNA was detected in all three sample replicates at only 2 of 8 sites where larvae were observed (Table S6). This indicates that, as with most eDNA sampling, replication is needed to achieve reasonable detection probabilities in low density populations. For western spadefoot, we detected eDNA in all but one site where larvae were observed at the time of sampling. Our rate of false negatives was similar that of an eDNA study in *A. boreas* (Franklin et al. 2018). However, we did detect western spadefoot with eDNA at one site where it was not observed visually. It is important to emphasize that eDNA sampling does not provide perfect detection, and future studies should seek to conduct more systematic eDNA sampling for arroyo toad and western spadefoot to determine detection probability of our eDNA assays more precisely.

The occupied range and abundance of arroyo toads and western spadefoot has declined dramatically in the past century. Despite recent efforts to protect them, ongoing drought and climate change may continue to have a negative impact on both species given their specialized breeding ecologies. However, climate change may increase the frequency of El Niño Southern Oscillation events (Ying et al. 2022) which typically causes increased precipitation in southern CA and Baja California. This could mitigate the effects of drought if El Niño events become more frequent than amphibian lifespans. Regardless, efficient and economical tools are needed to aid in monitoring of both species for informed management, and to evaluate the outcomes of conservation actions taken to protect them. We believe the sensitive, species-specific eDNA assays we designed for detection of arroyo toad and western spadefoot will provide managers with an important tool to aid in the monitoring and conservation of these imperiled species alongside traditional survey methods.

## ACKNOWLEDGEMENTS

We thank the Department of Defense Strategic Environmental Research and Development program for funding this work. We thank many biologists from MCBCP for helping with validation field work and logistical support including Sherri Sullivan, Colin Lee, Damien Cie, James Asmus, and Kathryn Carmody. We thank Valerie Hubbard from the USFS for help with validation field work and logistics in the Los Padres NF. We thank Robert Fisher for providing data on USGS 2022 arroyo toad surveys. We thank John LeBonte for help with western spadefoot validation, and Steve Junak for allowing access to his ranch for sampling. We thank Samuel Sweet for sharing his arroyo toad expertise. We thank Lauren Scheinberg from the California Academy of Sciences, Carol Spencer from the Berkeley Museum of Vertebrate Zoology, and Tara Luckau from UCLA for preparing and sending tissue samples.

## LITERATURE CITED

Carim, K. J. McKelvey, K. S. Young, M. K. Wilcox T. M., and M. K. Schwartz. 2016. A Protocol for Collecting Environmental DNA Samples From Streams. US Forset Service Rocky Mountain Research Station, General Technical Report RMRS-GTR-355

Fediajevaite, J., V. Priestley, R. Arnold, and V. Savolainen. 2021. Meta-analysis shows that environmental DNA outperforms traditional surveys, but warrants better reporting standards. Ecology and Evolution 11:4803–4815.

Fisher, R. N., and H. B. Shaffer. 1996. The Decline of Amphibians in California’s Great Central Valley. Conservation Biology 10:1387–1397.

Franklin, T. W., J. C. Dysthe, M. Golden, K. S. McKelvey, B. R. Hossack, K. J. Carim, C. Tait, M. K. Young, and M. K. Schwartz. 2018. Inferring presence of the western toad (Anaxyrus boreas) species complex using environmental DNA. Global Ecology and Conservation 15:e00438.

Gergus, E. W. A. 1998. Systematics of the Bufo microscaphus complex: allozyme evidence. Herpetologica 54:317–325.

Goldberg, C. S., C. R. Turner, K. Deiner, K. E. Klymus, P. F. Thomsen, M. A. Murphy, S. F. Spear, A. McKee, S. J. Oyler-McCance, R. S. Cornman, M. B. Laramie, A. R. Mahon, R. F. Lance, D. S. Pilliod, K. M. Strickler, L. P. Waits, A. K. Fremier, T. Takahara, J. E. Herder, and P. Taberlet. 2016. Critical considerations for the application of environmental DNA methods to detect aquatic species. Methods in Ecology and Evolution 7:1299–1307.

Hitchcock, C. J., E. A. Gallegos, A. R. Backlin, R. Barabe, P. H. Bloom, K. Boss, C. S. Brehme, C. W. Brown, D. R. Clark, E. R. Clark, K. Cooper, J. Donnell, E. Ervin, P. Famolaro, K. M. Guilliam, J. J. Hancock, N. Hess, S. Howard, V. Hubbartt, P. Lieske, R. Lovich, T. Matsuda, K. Meyer-Wilkins, K. Muri, B. Nerhus, J. Nordland, B. Ortega, R. Packard, R. Ramirez, S. C. Stewart, S. Sweet, M. Warburton, J. Wells, R. Winkleman, K. Winter, B. Zitt, and R. N. Fisher. 2022. Range-wide persistence of the endangered arroyo toad (Anaxyrus californicus) for 20+ years following a prolonged drought. Ecology and Evolution 12:e8796.

Jennings, M., and M. Hayes. 1994. Amphibian and Reptile Species of Special Concern in California. California Department of Fish and Game special report, contract number 8023

Neal, K. M., R. N. Fisher, M. J. Mitrovich, and H. B. Shaffer. 2020. Conservation Genomics of the Threatened Western Spadefoot, Spea hammondii, in Urbanized Southern California. Journal of Heredity 111:613–627.

Neal, K. M., B. B. Johnson, and H. B. Shaffer. 2018. Genetic structure and environmental niche modeling confirm two evolutionary and conservation units within the western spadefoot (Spea hammondii). Conservation Genetics 19:937–946.

Thomson R. C., A. N. Wright., and H. B. Shaffer H. B. 2016. California amphibian and reptile species of special concern. UC Press, Berkeley.

USFWS. 2014. Arroyo toad (Anaxyrus californicus) species report.

Ventura, CA. Wilcox, T. M., K. S. McKelvey, M. K. Young, S. F. Jane, W. H. Lowe, A. R. Whiteley, and M. K. Schwartz. 2013. Robust Detection of Rare Species Using Environmental DNA: The Importance of Primer Specificity. PLoS ONE 8.

Wright, E. S., L. S. Yilmaz, S. Ram, J. M. Gasser, G. W. Harrington, and D. R. Noguera. 2014. Exploiting extension bias in polymerase chain reaction to improve primer specificity in ensembles of nearly identical DNA templates. Environmental Microbiology 16:1354–1365.

Ye, J., G. Coulouris, I. Zaretskaya, I. Cutcutache, S. Rozen, and T. L. Madden. 2012. Primer-BLAST: a tool to design target-specific primers for polymerase chain reaction. BMC bioinformatics 13:134.

Ying, J., M. Collins, W. Cai, A. Timmermann, P. Huang, D. Chen, and K. Stein. 2022. Emergence of climate change in the tropical Pacific. Nature Climate Change 12:356–364.

