## Supplementary material for "Multiplex qPCR assays for detection of two imperiled anuran species, *Anaxyrus californicus* and *Spea hammondii*, from environmental DNA": Suppliment

***Supplementary Information***

**Table S1.** *Anaxyrus californicus* tissue samples sequenced for design of a species-specific qPCR assay for detection from environmental DNA. CAS is California Academy of Sciences, MVZ is Museum of Vertebrate Zoology.

| **Sample name** | **Country** | **County** | **Drainage** | **Date collected** | **Latitude** | **Longitude** | **Source** | **GenBank Accession #** |
| --- | --- | --- | --- | --- | --- | --- | --- | --- |
| RT97 | USA | San Diego | Kitchen Creek | 8/11/1990 | 32.7682 | -116.45098 | [CAS 175636](https://researcharchive.calacademy.org/research/herpetology/catalog/index.asp?xAction=getrec&close=true&CatalogNo=CAS+175636) | OP701659 |
| RT98 | USA | San Diego | Pine Valley Creek | 5/8/1991 | 32.84328 | -116.52676 | [CAS 178994](https://researcharchive.calacademy.org/research/herpetology/catalog/index.asp?xAction=getrec&close=true&CatalogNo=CAS+178994) | OP701679 |
| RT99 | USA | San Diego | San Luis Rey River | 5/7/1991 | 33.21301 | -116.56608 | [CAS 179041](https://researcharchive.calacademy.org/research/herpetology/catalog/index.asp?xAction=getrec&close=true&CatalogNo=CAS+179041) | OP701680 |
| RT162 | Mexico | NA | Santo Domingo River | 5/20/1977 | 30.75531 | -115.95714 | [MVZ:Herp:145230](http://arctos.database.museum/guid/MVZ:Herp:145230) | OP718187 |
| RT163 | Mexico | NA | Santo Domingo River | 5/20/1977 | 30.75531 | -115.95714 | [MVZ:Herp:150013](http://arctos.database.museum/guid/MVZ:Herp:150013) | OP718192 |
| RL1 | USA | Los Angeles | Castaic Creek | Unknown | Unknown | Unknown | Private collection | OP701646 |
| RL10 | USA | San Diego | Sweetwater River | 3/5/2001 | Unknown | Unknown | Private collection | OP701647 |
| RL11 | USA | San Diego | Sweetwater River | 3/5/2001 | Unknown | Unknown | Private collection | OP701660 |
| RL12 | USA | San Diego | Sweetwater River | 3/5/2001 | Unknown | Unknown | Private collection | OP701648 |
| RL13 | USA | San Diego | Sweetwater River | 3/5/2001 | Unknown | Unknown | Private collection | OP701649 |
| RL14 | USA | San Diego | Sweetwater River | 3/5/2001 | Unknown | Unknown | Private collection | OP701650 |
| RL15 | USA | San Diego | Sweetwater River | 3/5/2001 | Unknown | Unknown | Private collection | OP701651 |
| RL19 | USA | San Diego | San Luis Rey River | 4/14/2001 | 33.33204 | -117.15096 | Private collection | OP701661 |
| RL20 | USA | San Diego | San Luis Rey River | 4/14/2001 | 33.33202 | -117.15083 | Private collection | OP701652 |
| RL21 | USA | San Diego | Santa Ysabel Creek | 4/17/2001 | 33.11761 | -116.89404 | Private collection | OP701671 |
| RL22 | USA | San Diego | Santa Ysabel Creek | 4/17/2001 | 33.11945 | -116.89501 | Private collection | OP701672 |
| RL23 | USA | San Diego | Santa Ysabel Creek | 4/17/2001 | 33.11905 | -116.89494 | Private collection | OP701673 |
| RL24 | USA | San Diego | Santa Ysabel Creek | 4/17/2001 | 33.09236 | -116.89591 | Private collection | OP701674 |
| RL25 | USA | San Diego | Santa Ysabel Creek | 4/17/2001 | 33.11774 | -116.89404 | Private collection | OP718188 |
| RL26 | USA | San Diego | Santa Ysabel Creek | 4/17/2001 | 33.09216 | -116.89604 | Private collection | OP701662 |
| RL27 | USA | Orange | Gabino Creek | 4/19/2001 | 33.47949 | -117.53974 | Private collection | OP718177 |
| RL28 | USA | Orange | Gabino Creek | 4/19/2001 | 33.47965 | -117.5397 | Private collection | OP718178 |
| RL29 | USA | San Diego | DeLuz Creek | 4/21/2001 | 33.4153 | -117.31903 | Private collection | OP701653 |
| RL3 | USA | San Diego | San Mateo Creek | 12/7/2000 | 33.4683 | -117.47966 | Private collection | OP718179 |
| RL31 | USA | San Bernardino | Mojave River | 5/9/2001 | 34.34094 | -117.2453 | Private collection | OP718180 |
| RL32 | USA | Riverside | Santa Rosa Plateau | 5/9/2001 | 33.5443 | -117.2782 | Private collection | OP718189 |
| RL35 | USA | Monterrey | San Antonio Creek | 5/21/2001 | 35.99911 | -121.24664 | Private collection | OP701629 |
| RL36 | USA | Monterrey | San Antonio Creek | 5/21/2001 | 36.00035 | -121.24805 | Private collection | OP701630 |
| RL37 | USA | Monterrey | San Antonio Creek | 5/21/2001 | 35.99803 | -121.24688 | Private collection | OP701631 |
| RL38 | USA | Monterrey | San Antonio Creek | 5/21/2001 | 35.91269 | -121.13388 | Private collection | OP701632 |
| RL39 | USA | Monterrey | San Antonio Creek | 5/21/2001 | 35.91269 | -121.13388 | Private collection | OP701633 |
| RL4 | USA | San Diego | San Mateo Creek | 12/11/2000 | 33.46888 | -117.47929 | Private collection | OP701654 |
| RL41 | USA | Santa Barbara | Sata Ynez River | 5/22/2001 | 34.51316 | -119.59407 | Private collection | OP701634 |
| RL42 | USA | Santa Barbara | Mono Creek | 5/22/2001 | 34.53566 | -119.63026 | Private collection | OP701635 |
| RL46 | USA | Santa Barbara | Indian Creek | 5/22/2001 | 34.53655 | -119.63342 | Private collection | OP701636 |
| RL49 | USA | Ventura | Piru Creek | 5/23/2001 | 34.69102 | -118.85123 | Private collection | OP718182 |
| RL5 | USA | San Diego | San Mateo Creek | 4/20/2001 | 33.46888 | -117.47929 | Private collection | OP701655 |
| RL54 | Mexico | NA | La Mision | 6/8/2001 | 32.10069 | -116.80988 | Private collection | OP718184 |
| RL56 | Mexico | NA | La Mision | 6/8/2001 | 32.10066 | -116.8099 | Private collection | OP718185 |
| RL57 | Mexico | NA | La Mision | 6/8/2001 | 32.10066 | -116.8099 | Private collection | OP701663 |
| RL6 | USA | San Diego | Santa Ysabel Creek | 1/10/2000 | 33.08513 | -116.98833 | Private collection | OP701664 |
| RL62 | USA | Riverside | Bautista Creek | 6/22/2001 | 33.64867 | -116.81616 | Private collection | OP701656 |
| RL64 | USA | Los Angeles | Littlerock Creek | 6/30/2001 | 34.46017 | -118.0194 | Private collection | OP701637 |
| RL65 | USA | Los Angeles | Littlerock Creek | 6/30/2001 | 34.45898 | -118.01861 | Private collection | OP701638 |
| RL66 | USA | Los Angeles | Tujunga Creek | 6/30/2001 | 34.31051 | -118.09439 | Private collection | OP701639 |
| RL67 | Mexico | NA | Guadalupe | 3/22/2002 | 32.11827 | -116.47655 | Private collection | OP701665 |
| RL68 | Mexico | NA | Guadalupe | 3/22/2002 | 32.11803 | -116.47668 | Private collection | OP701666 |
| RL69 | Mexico | NA | Las Palmas | 3/23/2002 | 32.3303 | -116.51396 | Private collection | OP718190 |
| RL7 | USA | San Diego | Santa Ysabel Creek | 1/10/2000 | 33.08525 | -116.98805 | Private collection | OP701667 |
| RL70 | Mexico | NA | Maeadero | 4/5/2002 | 31.63665 | -116.4657 | Private collection | OP718171 |
| RL71 | Mexico | NA | San Vicete | 5/17/2002 | 31.33415 | -116.4657 | Private collection | OP718172 |
| RL72 | Mexico | NA | San Vicete | 5/17/2002 | 31.33415 | -116.4657 | Private collection | OP701675 |
| RL73 | Mexico | NA | San Vicete | 5/17/2002 | 31.33415 | -116.4657 | Private collection | OP701676 |
| RL75 | Mexico | NA | San Rafael | 5/25/2002 | 31.09876 | -116.02539 | Private collection | OP718173 |
| RL76 | Mexico | NA | San Rafael | 5/25/2002 | 31.09876 | -116.02539 | Private collection | OP701677 |
| RL8 | USA | San Diego | Cottonwood Creek | 3/2/2001 | 32.57298 | -116.75718 | Private collection | OP701668 |
| RL87 | USA | San Diego | San Diego River | Unknown | Unknown | Unknown | Private collection | OP701669 |
| RL88 | USA | San Diego | San Diego River | Unknown | Unknown | Unknown | Private collection | OP701670 |
| RL89 | Mexico | NA | Santo Tomas | 5/26/2004 | 31.54913 | -116.22174 | Private collection | OP718174 |
| RL9 | USA | San Diego | Sweetwater River | 3/5/2001 | Unknown | Unknown | Private collection | OP701657 |
| RL90 | Mexico | NA | Santo Tomas | 5/26/2004 | 31.54913 | -116.22174 | Private collection | OP718175 |
| RL91 | Mexico | NA | Santo Tomas | 5/26/2004 | 31.54913 | -116.22174 | Private collection | OP718176 |
| RL93 | Mexico | NA | Rio San Telmo | 4/14/2005 | 31.9775 | -115.74525 | Private collection | OP718186 |
| RL94 | Mexico | NA | Rio Santa Maria | 4/15/2005 | 31.51107 | -115.80393 | Private collection | OP718191 |
| RT185 (RL2) | USA | San Diego | Tecate Creek | 5/18/1999 | 33.51444 | -117.57166 | Private collection | OP701658 |
| RT186 (RL18) | USA | San Diego | Cottonwood Creek | 3/29/2001 | 32.57368 | -116.7565 | Private collection | OP701640 |
| RT187 (RL34) | USA | San Bernardino | Mojave River | 5/17/2001 | 34.32394 | -117.40901 | Private collection | OP718181 |
| RT190 (RL40) | USA | Monterrey | San Antonio Creek | 5/21/2001 | 35.91269 | -121.13388 | Private collection | OP701641 |
| RT191 (RL43) | USA | Santa Barbara | Mono Creek | 5/22/2001 | 34.53665 | -119.63215 | Private collection | OP701642 |
| RT192 (RL45) | USA | Santa Barbara | Mono Creek | 5/22/2001 | 34.53665 | -119.63215 | Private collection | OP701643 |
| RT193 (RL47) | USA | Santa Barbara | Indian Creek | 5/22/2001 | 34.54031 | -119.63641 | Private collection | OP701644 |
| RT194 (RL50) | USA | Ventura | Piru Creek | 5/23/2001 | 34.69199 | -188.85308 | Private collection | OP718183 |
| RT196 (RL63) | USA | Santa Barbara | Sisquoc River | 6/29/2001 | 34.827 | -119.99904 | Private collection | OP701645 |
| RT198 (RL106) | USA | San Diego | Pine Valley Creek | 5/5/2008 | 32.85335 | -116.52256 | Private collection | OP701678 |

**Table S2.** *Spea hammondii* tissue samples sequenced for design of a species-specific qPCR assay for detection from environmental DNA. CAS is California Academy of Sciences, MVZ is Museum of Vertebrate Zoology, and HBS is private collection of H. Bradley Shaffer.

| **Sample name** |  | **County** | **Date collected** | **Latitude** | **Longitude** | **Source** | **GenBank Accession #** | |
| --- | --- | --- | --- | --- | --- | --- | --- | --- |
|  | **Country** |  |  |  |  |  | **cytb** | **16S** |
| RT125 | USA | Tulare | 5/19/2001 | 35.89375 | -118.9088 | [CAS-223542](http://researcharchive.calacademy.org/research/herpetology/catalog/index.asp?xAction=getrec&close=true&CatalogNo=CAS+223542) | OP265969 | MT137273 |
| RT129 | USA | Stanislaus | 5/20/2001 | 37.45318 | -121.27483 | [CAS-225300](http://researcharchive.calacademy.org/research/herpetology/catalog/index.asp?xAction=getrec&close=true&CatalogNo=CAS+225300) | OP265970 | OP131912 |
| RT166 | USA | San Joaquin | 11/17/1996 | 37.60078 | -121.53529 | [MVZ:Herp:234174](http://arctos.database.museum/guid/MVZ:Herp:234174) | OP265971 | OP131913 |
| RT167 | USA | San Joaquin | 1/15/1997 | 37.67221 | -121.45633 | [MVZ:Herp:234175](http://arctos.database.museum/guid/MVZ:Herp:234175) | OP265972 | MT135561 |
| RT199 | USA | Glenn | 4/7/1990 | 39.79739 | -122.55138 | HBS-9764 | OP227147 | not Sequenced |
| RT200 | USA | Glenn | 4/7/1990 | 39.79739 | -122.55138 | HBS-9765 | OP227158 | OP115847 |
| RT201 | USA | Yolo | 5/16/1998 | 38.91153 | -122.02286 | HBS-22298 | OP227169 | not Sequenced |
| RT202 | USA | Yolo | 5/16/1998 | 38.91153 | -122.02286 | HBS-22299 | OP227179 | not Sequenced |
| RT203 | USA | San Luis Obispo | 4/28/2017 | 35.1569 | -119.96882 | HBS-132798 | OP227180 | OP115848 |
| RT204 | USA | San Luis Obispo | 4/28/2017 | 35.1569 | -119.96882 | HBS-132799 | OP227181 | OP115849 |
| RT205 | USA | San Luis Obispo | 4/28/2017 | 35.1569 | -119.96882 | HBS-132800 | OP227182 | OP115850 |
| RT206 | USA | San Luis Obispo | 4/28/2017 | 35.1569 | -119.96882 | HBS-132801 | OP227183 | OP115851 |
| RT207 | USA | San Luis Obispo | 4/28/2017 | 35.1569 | -119.96882 | HBS-132802 | OP227184 | OP115852 |
| RT208 | USA | San Luis Obispo | 4/14/2017 | 35.7821 | -120.33481 | HBS-133582 | OP227148 | OP115853 |
| RT209 | USA | San Luis Obispo | 4/14/2017 | 35.7821 | -120.33481 | HBS-133583 | OP227149 | OP115854 |
| RT210 | USA | San Luis Obispo | 4/14/2017 | 35.7821 | -120.33481 | HBS-133584 | OP227150 | OP115855 |
| RT211 | USA | San Luis Obispo | 4/14/2017 | 35.7821 | -120.33481 | HBS-133585 | OP227151 | OP115856 |
| RT213 | USA | Santa Barbara | 5/3/2018 | 34.768143 | -120.393662 | HBS-134538 | OP227152 | OP115857 |
| RT214 | USA | Santa Barbara | 5/3/2018 | 34.768143 | -120.393662 | HBS-134539 | OP227153 | OP115858 |
| RT215 | USA | Santa Barbara | 5/3/2018 | 34.768143 | -120.393662 | HBS-134540 | OP227154 | OP115859 |
| RT216 | USA | Santa Barbara | 5/3/2018 | 34.768143 | -120.393662 | HBS-134541 | OP227155 | OP115860 |
| RT217 | USA | Santa Barbara | 5/3/2018 | 34.768143 | -120.393662 | HBS-134542 | OP227156 | OP115861 |
| RT218 | USA | Kern | 3/18/2017 | 35.03829 | -118.71936 | HBS-131797 | OP227157 | OP115862 |
| RT219 | USA | Kern | 3/18/2017 | 35.03829 | -118.71936 | HBS-131799 | OP227159 | OP115863 |
| RT220 | USA | Orange | 2/2/2017 | 33.5635 | -117.80741 | HBS-132561 | OP227160 | OP115732 |
| RT221 | USA | Orange | 2/2/2017 | 33.5635 | -117.80741 | HBS-132563 | OP227161 | OP115733 |
| RT222 | USA | Orange | 3/21/2017 | 33.60307 | -117.55997 | HBS-132145 | OP227162 | OP115734 |
| RT223 | USA | Orange | 3/21/2017 | 33.60307 | -117.55997 | HBS-132146 | OP227163 | OP115735 |
| RT224 | USA | Ventura | 3/5/2017 | 34.32077 | -118.86572 | HBS-131710 | OP227164 | OP115736 |
| RT225 | USA | Ventura | 3/5/2017 | 34.32077 | -118.86572 | HBS-131712 | OP227165 | OP115737 |
| RT226 | USA | San Diego | 4/3/2017 | 32.7125 | -116.916 | HBS-131669 | OP227166 | OP115738 |
| RT227 | USA | San Diego | 4/3/2017 | 32.7125 | -116.916 | HBS-131670 | OP227167 | OP115739 |
| RT228 | USA | Monterey | 4/10/2017 | 36.50333 | -121.34906 | HBS-133222 | OP227168 | OP115864 |
| RT229 | USA | Monterey | 4/10/2017 | 36.50333 | -121.34906 | HBS-133223 | OP227170 | OP115865 |
| RT230 | USA | Monterey | 4/14/2017 | 35.83887 | -120.38189 | HBS-133556 | OP227171 | OP115866 |
| RT231 | USA | Monterey | 4/14/2017 | 35.83887 | -120.38189 | HBS-133557 | OP227172 | OP115867 |
| RT232 | USA | Monterey | 4/14/2017 | 35.83887 | -120.38189 | HBS-133558 | OP227173 | OP115868 |
| RT233 | USA | Monterey | 4/14/2017 | 35.83887 | -120.38189 | HBS-133568 | OP227174 | OP115869 |
| RT234 | USA | Monterey | 4/14/2017 | 35.83887 | -120.38189 | HBS-133569 | OP227175 | OP115870 |
| RT235 | USA | Monterey | 4/14/2017 | 35.83887 | -120.38189 | HBS-133570 | OP227176 | OP115871 |
| RT236 | USA | Monterey | 4/14/2017 | 35.83887 | -120.38189 | HBS-133571 | OP227177 | OP115872 |
| RT237 | USA | Monterey | 4/14/2017 | 35.83887 | -120.38189 | HBS-133572 | OP227178 | OP115873 |

**Table S3.** Non-target species tissue samples sequenced for design of species-specific qPCR assays for detection of *Anaxyrus californicus* and *Spea hammondii* from environmental DNA.

| **Sample name** | **Species** | **Common name** | **State** | **County** | **Source** | **GenBank Accession #** | |
| --- | --- | --- | --- | --- | --- | --- | --- |
|  |  |  |  |  |  | **Cytb** | **16s** |
| RT100 | *Rana draytonii* | California red-legged frog | CA | Contra Costa | [CAS-200288](http://researcharchive.calacademy.org/research/herpetology/catalog/index.asp?xAction=getrec&close=true&CatalogNo=CAS+200288) | OP391557 | MT137263 |
| RT102 | *Rana draytonii* | California red-legged frog | CA | Contra Costa | [CAS-200290](http://researcharchive.calacademy.org/research/herpetology/catalog/index.asp?xAction=getrec&close=true&CatalogNo=CAS+200290) | OP391558 | MT137264 |
| RT103 | *Ensatina eschscholtzii xanthoptica* | Yellow-eyed Ensatina salamander | CA | Santa Cruz | [CAS-203543](http://researcharchive.calacademy.org/research/herpetology/catalog/index.asp?xAction=getrec&close=true&CatalogNo=CAS+203543) | OP391538 | not sequenced |
| RT105 | *Rana boylii* | Foothill yellow-legged frog | CA | Santa Clara | [CAS-205752](http://researcharchive.calacademy.org/research/herpetology/catalog/index.asp?xAction=getrec&close=true&CatalogNo=CAS+205752) | OP391559 | MT137265 |
| RT106 | *Ensatina eschscholtzii eschscholtzii* | Ensatina salamander | CA | Monterey | [CAS-205796](http://researcharchive.calacademy.org/research/herpetology/catalog/index.asp?xAction=getrec&close=true&CatalogNo=CAS+205796) | OP391540 | MT137266 |
| RT108 | *Anaxyrus boreas halophilus* | Western toad | CA | Monterey | [CAS-206465](http://researcharchive.calacademy.org/research/herpetology/catalog/index.asp?xAction=getrec&close=true&CatalogNo=CAS+206465) | OP391560 | MT137267 |
| RT109 | *Ensatina eschscholtzii xanthoptica* | Yellow-eyed Ensatina salamander | CA | San Mateo | [CAS-207429](http://researcharchive.calacademy.org/research/herpetology/catalog/index.asp?xAction=getrec&close=true&CatalogNo=CAS+207429) | OP391541 | MT137268 |
| RT111 | *Pseudacris sierra* | Sierran chorus frog | CA | San Luis Obispo | [CAS-208510](http://researcharchive.calacademy.org/research/herpetology/catalog/index.asp?xAction=getrec&close=true&CatalogNo=CAS+208510) | not sequenced | MT137269 |
| RT112 | *Aneides lugubris* | Arboreal salamander | CA | Santa Cruz | [CAS-208660](http://researcharchive.calacademy.org/research/herpetology/catalog/index.asp?xAction=getrec&close=true&CatalogNo=CAS+208660) | OP391542 | MT137270 |
| RT118 | *Sternotherus odoratus* | Common musk turtle | FL | Taylor | [CAS-214343](http://researcharchive.calacademy.org/research/herpetology/catalog/index.asp?xAction=getrec&close=true&CatalogNo=CAS+214343) | not sequenced | MT137271 |
| RT120 | *Batrachoseps nigriventris* | Black-bellied slender salamander | CA | San Luis Obispo | [CAS-214856](http://researcharchive.calacademy.org/research/herpetology/catalog/index.asp?xAction=getrec&close=true&CatalogNo=CAS+214856) | OP391543 | MT137272 |
| RT121 | *Aneides lugubris* | Arboreal salamander | CA | Monterey | [CAS-214883](http://researcharchive.calacademy.org/research/herpetology/catalog/index.asp?xAction=getrec&close=true&CatalogNo=CAS+214883) | OP391544 | not sequenced |
| RT123 | *Ambystoma californiense* | California Tiger Salamander | CA | Monterey | [CAS-214885](http://researcharchive.calacademy.org/research/herpetology/catalog/index.asp?xAction=getrec&close=true&CatalogNo=CAS+214885) | OP391551 | not sequenced |
| RT127 | *Lithobates catesbeiana* | America Bullfrog | CA | Fresno | [CAS-224776](http://researcharchive.calacademy.org/research/herpetology/catalog/index.asp?xAction=getrec&close=true&CatalogNo=CAS+224776) | OP391561 | MT137274 |
| RT131 | *Rana boylii* | Foothill yellow-legged frog | CA | Santa Clara | [CAS-226108](http://researcharchive.calacademy.org/research/herpetology/catalog/index.asp?xAction=getrec&close=true&CatalogNo=CAS+226108) | OP391562 | MT137275 |
| RT135 | *Trachemys scripta* | Pond slider | GA | Shasta | [CAS-227634](http://researcharchive.calacademy.org/research/herpetology/catalog/index.asp?xAction=getrec&close=true&CatalogNo=CAS+227634) | not sequenced | MT137276 |
| RT142 | *Xenopus laevis* | African clawed frog | CA | San Francisco | [CAS-244034](http://researcharchive.calacademy.org/research/herpetology/catalog/index.asp?xAction=getrec&close=true&CatalogNo=CAS+244034) | OP391555 | MT137278 |
| RT145 | *Batrachoseps luciae* | Santa Lucia Mountains slender slamander | CA | Monterey | [CAS-252904](http://researcharchive.calacademy.org/research/herpetology/catalog/index.asp?xAction=getrec&close=true&CatalogNo=CAS+252904) | OP391545 | MT137279 |
| RT146 | *Batrachoseps luciae* | Santa Lucia Mountains slender slamander | CA | Monterey | [CAS-252906](http://researcharchive.calacademy.org/research/herpetology/catalog/index.asp?xAction=getrec&close=true&CatalogNo=CAS+252906) | OP391546 | not sequenced |
| RT147 | *Pseudacris sierra* | Sierran chorus frog | CA | Monterey | [CAS-252908](http://researcharchive.calacademy.org/research/herpetology/catalog/Index.asp) | OP391563 | not sequenced |
| RT148 | *Ensatina eschscholtzii* | Ensatina salamander | CA | Monterey | [CAS-252913](http://researcharchive.calacademy.org/research/herpetology/catalog/index.asp?xAction=getrec&close=true&CatalogNo=CAS+252913) | OP391547 | not sequenced |
| RT150 | *Anaxyrus boreas halophilus* | Western Toad | CA | Kern | [CAS-253026](http://researcharchive.calacademy.org/research/herpetology/catalog/index.asp?xAction=getrec&close=true&CatalogNo=CAS+253026) | OP391564 | MT137280 |
| RT151 | *Lithobates catesbeiana* | America Bullfrog | CA | Kern | [CAS-253042](http://researcharchive.calacademy.org/research/herpetology/catalog/index.asp?xAction=getrec&close=true&CatalogNo=CAS+253042) | OP391565 | MT137281 |
| RT152 | *Taricha torosa* | California newt | GA | Kern | [CAS-253044](http://researcharchive.calacademy.org/research/herpetology/catalog/index.asp?xAction=getrec&close=true&CatalogNo=CAS+253044) | OP391552 | OP378122 |
| RT153 | *Xenopus laevis* | African clawed frog | CA | San Francisco | [CAS-253186](http://researcharchive.calacademy.org/research/herpetology/catalog/index.asp?xAction=getrec&close=true&CatalogNo=CAS+253186) | OP391556 | MT137282 |
| RT154 | *Sternotherus odoratus* | Common musk turtle | GA | Macon | [CAS-255136](http://researcharchive.calacademy.org/research/herpetology/catalog/index.asp?xAction=getrec&close=true&CatalogNo=CAS+255136) | not sequenced | MT137283 |
| RT155 | *Pseudacris cadaverina* | California chorus frog | CA | Los Angeles | [CAS-255413](http://researcharchive.calacademy.org/research/herpetology/catalog/index.asp?xAction=getrec&close=true&CatalogNo=CAS+255413) | OP391566 | MT137284 |
| RT159 | *Taricha torosa sierrae* | Sierra newt | GA | Madera | [CAS-259608](http://researcharchive.calacademy.org/research/herpetology/catalog/index.asp?xAction=getrec&close=true&CatalogNo=CAS+259608) | OP391553 | OP378123 |
| RT160 | *Rana boylii* | Foothill yellow-legged frog | CA | Amador | [CAS-259672](http://researcharchive.calacademy.org/research/herpetology/catalog/index.asp?xAction=getrec&close=true&CatalogNo=CAS+259672) | OP391567 | MT137285 |
| RT165 | *Pseudacris cadaverina* | California chorus frog | Mexico | Baja Norte | [MVZ:Herp:145382](http://arctos.database.museum/guid/MVZ:Herp:145382) | OP391568 | MT135560 |
| RT168 | *Batrachoseps incognitus* | San Simeon slender salamander | CA | Monterey | [MVZ:Herp:251931](http://arctos.database.museum/guid/MVZ:Herp:251931) | OP391539 | MT135562 |
| RT171 | *Taricha torosa* | California newt | GA | San Luis Obispo | [MVZ:Herp:236242](http://arctos.database.museum/guid/MVZ:Herp:236242) | OP391554 | OP378124 |
| RT173 | *Actinemys marmorata* | Western pond turtle | CA | Napa | [MVZ:Herp:164994](http://arctos.database.museum/guid/MVZ:Herp:164994) | not sequenced | MT135564 |
| RT174 | *Actinemys marmorata* | Western pond turtle | CA | Lake | [MVZ:Herp:164995](http://arctos.database.museum/guid/MVZ:Herp:164995) | not sequenced | MT135565 |
| RT175 | *Chrysemys picta* | Painted turtle | NM | Sierra | [MVZ:Herp:250712](http://arctos.database.museum/guid/MVZ:Herp:250712) | not sequenced | MT135566 |
| RT176 | *Chrysemys picta* | Painted turtle | WA | Spokane | [MVZ:Herp:238581](http://arctos.database.museum/guid/MVZ:Herp:238581) | not sequenced | MT135567 |
| RT177 | *Trachemys scripta elegans* | Red-eared slider | NM | Sierra | [MVZ:Herp:265667](http://arctos.database.museum/guid/MVZ:Herp:265667) | not sequenced | MT135568 |
| RT238 | *Pseudacris hypochondriaca* | Baja California chorus frog | CA | Riverside | [CAS-200610](http://researcharchive.calacademy.org/research/herpetology/catalog/index.asp?xAction=getrec&close=true&CatalogNo=CAS+200610) | not sequenced | OP377739 |
| RT239 | *Pseudacris hypochondriaca* | Baja California chorus frog | CA | Riverside | [CAS-200616](http://researcharchive.calacademy.org/research/herpetology/catalog/index.asp?xAction=getrec&close=true&CatalogNo=CAS+200616) | not sequenced | OP377740 |
| RT87 | *Ambystoma californiense* | California tiger salamander | CA | Santa Clara | [CAS-211685](http://researcharchive.calacademy.org/research/herpetology/catalog/index.asp?xAction=getrec&close=true&CatalogNo=CAS+211685) | OP391548 | not sequenced |
| RT93 | *Ambystoma californiense* | California tiger salamander | CA | Contra Costa | [CAS-238956](http://researcharchive.calacademy.org/research/herpetology/catalog/index.asp?xAction=getrec&close=true&CatalogNo=CAS+238956) | OP391549 | not sequenced |
| RT95 | *Taricha torosa sierrae* | Sierra newt | CA | Fresno | [CAS-208714](http://researcharchive.calacademy.org/research/herpetology/catalog/index.asp?xAction=getrec&close=true&CatalogNo=CAS+208714) | OP391550 | OP378125 |

**Table S4.** Sequences obtained from GenBank used in design of species-specific qPCR assays for detection of *Anaxyrus californicus* and *Spea hammondii* from environmental DNA.

| **Species Name** | **Common name** | **Gene** | **GenBank Accession #** |
| --- | --- | --- | --- |
| *Actinemys marmorata* | Western pond turtle | cytb | EU787059 |
| *Actinemys marmorata* | Western pond turtle | cytb | EU787064 |
| *Batrachoseps gavilanensis* | Gabilan Mountains slender salamander | cytb | JQ250323 |
| *Batrachoseps gavilanensis* | Gabilan Mountains slender salamander | cytb | JQ250324 |
| *Batrachoseps incognitus* | San Simeon slender salamander | cytb | JQ250328 |
| *Chrysemys picta* | Painted turtle | cytb | AF069423 |
| *Chrysemys picta* | Painted turtle | cytb | KJ536200 |
| *Pseudacris hypochondriaca* | Baja California chorus frog | cytb | KJ536199 |
| *Pseudacris hypochondriaca* | Baja California chorus frog | cytb | KJ536200 |
| *Pseudacris sierra* | Sierran chorus frog | cytb | KJ536202 |
| *Spea hammondii* | Western spadefoot | cytb | AY236788 |
| *Sternotherus odoratus* | Common musk turtle | cytb | MG460002 |
| *Sternotherus odoratus* | Common musk turtle | cytb | MG460122 |
| *Trachemys scripta elegans* | Red-eared slider | cytb | EU787024.1 |
| *Trachemys scripta elegans* | Red-eared slider | cytb | FJ770617.1 |
| *Ambystoma calforniense* | California tiger salamander | 16S | AY659995.1 |
| *Ambystoma tigrinum* | tiger salamander | 16S | AY659992.1 |
| *Ambystoma tigrinum* | tiger salamander | 16S | DQ283407.1 |
| *Anaxyrus boreas* | Western toad | 16S | AY680242.1 |
| *Anaxyrus boreas* | Western Toad | 16S | AY680244.1 |
| *Anaxyrus boreas* | Western toad | 16S | DQ158436.1 |
| *Anaxyrus boreas* | Western Toad | 16S | DQ283180.1 |
| *Anaxyrus boreas* | Western toad | 16S | KF665480.1 |
| *Anaxyrus boreas* | Western Toad | 16S | U52752.1 |
| *Aneides lugubris* | Arboreal salamander | 16S | DQ105346.1 |
| *Aneides lugubris* | Arboreal salamander | 16S | DQ105347.1 |
| *Aneides lugubris* | Arboreal salamander | 16S | DQ105348.1 |
| *Aneides lugubris* | Arboreal salamander | 16S | DQ105349.1 |
| *Aneides lugubris* | Arboreal salamander | 16S | DQ105350.1 |
| *Batrachoseps attenuatus* | California slender salamander | 16S | EU011483.1 |
| *Batrachoseps bramei* | Fairview slender salamander | 16S | JQ035722.1 |
| *Batrachoseps diabolicus* | Hell Hollow slender salamander | 16S | EU011256.1 |
| *Batrachoseps gabrieli* | San Gabriel slender salamander | 16S | AF199234.1 |
| *Batrachoseps gavilanensis* | Gabilan Mountains slender salamander | 16S | EU011260.1 |
| *Batrachoseps gregarius* | Gregarious slender salamander | 16S | JQ035727.1 |
| *Batrachoseps major* | Garden slender salamander | 16S | DQ642047.1 |
| *Batrachoseps nigriventris* | Black-bellied slender salamander | 16S | JQ035712.1 |
| *Batrachoseps nigriventris* | Black-bellied slender salamander | 16S | KT368149.1 |
| *Batrachoseps simatus* | Kern Canyon slender salamander | 16S | JQ035713.1 |
| *Batrachoseps stebbinsi* | Tehachapi slender salamander | 16S | JQ035728.1 |
| *Chrysemys picta* | Painted turtle | 16S | AF069423.1 |
| *Chrysemys picta* | Painted turtle | 16S | KF874616.1 |
| *Chrysemys picta* | Painted turtle | 16S | MN135379.1 |
| *Lithobates catesbeiana* | America Bullfrog | 16S | DQ283257.1 |
| *Lithobates catesbeiana* | America Bullfrog | 16S | KX269208.1 |
| *Lithobates catesbeiana* | America Bullfrog | 16S | KY677817.1 |
| *Lithobates catesbeiana* | America Bullfrog | 16S | M57527.1 |
| *Lithobates catesbeiana* | America Bullfrog | 16S | X12841.1 |
| *Pseudacris cadaverina* | America Bullfrog | 16S | AY291114.1 |
| *Pseudacris cadaverina* | America Bullfrog | 16S | AY843734.1 |
| *Pseudacris cadaverina* | America Bullfrog | 16S | EF472006.1 |
| *Pseudacris hypochondriaca* | Baja California chorus frog | 16S | AY291111.1 |
| *Pseudacris hypochondriaca* | Baja California chorus frog | 16S | AY843737.1 |
| *Pseudacris hypochondriaca* | Baja California chorus frog | 16S | DQ679375.1 |
| *Pseudacris hypochondriaca* | Baja California chorus frog | 16S | EF472005.1 |
| *Pseudacris hypochondriaca* | Baja California chorus frog | 16S | KY677793.1 |
| *Pseudacris hypochondriaca* | Baja California chorus frog | 16S | KY677803.1 |
| *Pseudacris regilla* | Pacific chorus frog | 16S | KY677809.1 |
| *Pseudacris sierra* | Sierran chorus frog | 16S | AY291112.1 |
| *Rana boylii* | Foothill yellow-legged frog | 16S | AY779192.1 |
| *Rana boylii* | Foothill yellow-legged frog | 16S | DQ347335.1 |
| *Rana draytonii* | California red-legged frog | 16S | DQ283189.1 |
| *Rana draytonii* | California red-legged frog | 16S | KP013110.1 |
| *Sternotherus odoratus* | Common musk turtle | 16S | KF301330.1 |
| *Sternotherus odoratus* | Common musk turtle | 16S | MH308386.1 |
| *Sternotherus odoratus* | Common musk turtle | 16S | MH308387.1 |
| *Sternotherus odoratus* | Common musk turtle | 16S | MH308388.1 |
| *Sternotherus odoratus* | Common musk turtle | 16S | MN135409.1 |
| *Trachemys scripta* | Pond slider | 16S | FJ392294.1 |
| *Trachemys scripta* | Pond slider | 16S | HQ123497.1 |
| *Trachemys scripta* | Pond slider | 16S | L28077.1 |
| *Trachemys scripta* | Pond slider | 16S | MN135556.1 |
| *Xenopus laevis* | African clawed frog | 16S | MH115794.1 |

**Table S5**. Candidate primers tested with SYBR Green qPCR for detection of *Anaxyrus californicus* and *Spea hammondii* from environmental DNA. Primers indicated in bold were ultimately chosen for the final analysis.

| **Primer/probe name** | **Sequence 5’-3’** | **Target gene** | **Tm (°C)** |
| --- | --- | --- | --- |
| ANCA1-F | GCTCAAATCGCAACTGGACTT | cytb | 58.1 |
| ANCA1-R | AATGAGGCTCCATTTGCATGT | cytb | 58.1 |
| ANCA2-F | AGCTGATACATCCATAGCCTTTTCAT | cytb | 59.4 |
| ANCA2-R | GAATGAGGCTCCATTTGCATGT | cytb | 59.9 |
| **ANCA3-F** | **TCACCAGCGAATATCTCCTCTTTAT** | cytb | 58.4 |
| **ANCA3-R** | **GGCTATGGATGTATCAGCTGTGTAA** | cytb | 58.3 |
| **SPHA1-F** | CGGAAACGTACTAGTCCAATGAATC | cytb | 59 |
| **SPHA1-R** | TCCGGTTTCGTGTAGGAATAGAAG | cytb | 59.5 |
| **SPHA2-F** | **CGGAAACGTACTAGTCCAATGAATC** | cytb | 59 |
| **SPHA2-R** | **GATCCGGTTTCGTGTAGGAATAGA** | cytb | 59.4 |
| SPHA3-F | TCTTAACCAAGAGCCACAGCTCTA | 16S | 58.4 |
| SPHA3-R | TGTCGATATGGACTCTTGAAATGG | 16S | 59 |
| SPHA4-F | CCCATGGAGCTTTAAACTTAAACC | 16S | 58.3 |
| SPHA4-R | CCCCAACCGAAAACATAAGC | 16S | 58 |

**Table S6.** eDNA samples collected for field validation of our species-specific qPCR assay for *Anaxyrus californicus.*

| **Sample name** | **Site Name** | **Water Body** | **Sample collection date** | **Latitude** | **Longitude** | ***A. californicus* visually observed** | ***A. californicus* detected with qPCR** |
| --- | --- | --- | --- | --- | --- | --- | --- |
| eDNA598 | Upper San Mateo sample 1 | San Mateo River | 5/8/2019 | 33.46897 | -117.47608 | Y | Y |
| eDNA599 | Upper San Mateo sample 2 | San Mateo River | 5/8/2019 | 33.41944 | -117.54919 | Y | Y |
| eDNA600 | Upper San Mateo sample 3 | San Mateo River | 5/8/2019 | 33.41944 | -117.54919 | Y | Y |
| eDNA601 | Christianitos sample 4 | Christianitos Creek | 5/8/2019 | 33.45113 | -117.56509 | Y | Y |
| eDNA602 | Lower San Onofre sample 5 | San Onofre River | 5/8/2019 | 33.39270 | -117.53036 | Y | Y |
| eDNA604 | Lower Margarita sample 6 | Santa Margarita River | 5/9/2019 | 33.28435 | -117.37415 | N | N |
| eDNA605 | Middle Margarita sample 7 | Santa Margarita River | 5/9/2019 | 33.32903 | -117.33542 | Y | Y |
| eDNA606 | Upper Margarita sample 8 | Santa Margarita River | 5/9/2019 | 33.36237 | -117.32151 | Y | Y |
| eDNA1694 | Agua Blanca-A | Agua Blanca Creek | 5/15/2021 | 34.54213 | -118.76582 | N | N |
| eDNA1695 | Agua Blanca-B | Agua Blanca Creek | 5/15/2021 | 34.54213 | -118.76582 | N | N |
| eDNA1696 | Agua Blanca-C | Agua Blanca Creek | 5/15/2021 | 34.54213 | -118.76582 | N | N |
| eDNA1697 | Mono-A | Mono Creek | 6/2/2021 | 34.54029 | -119.62549 | N | N |
| eDNA1698 | Mono-B | Mono Creek | 6/2/2021 | 34.54029 | -119.62549 | N | N |
| eDNA1699 | Mono-C | Mono Creek | 6/2/2021 | 34.54029 | -119.62549 | N | N |
| eDNA1700 | Indian-A | Indian Creek | 6/2/2021 | 34.54322 | -119.64208 | N | N |
| eDNA1701 | Indian-B | Indian Creek | 6/2/2021 | 34.54322 | -119.64208 | N | N |
| eDNA1702 | Indian-C | Indian Creek | 6/2/2021 | 34.54322 | -119.64208 | N | N |
| eDNA1703 | Santa Ynez-A | Santa Ynez River | 6/2/2021 | 34.51265 | -119.59246 | N | N |
| eDNA1704 | Santa Ynez-B | Santa Ynez River | 6/2/2021 | 34.51265 | -119.59246 | N | N |
| eDNA1705 | Santa Ynez-C | Santa Ynez River | 6/2/2021 | 34.51265 | -119.59246 | N | N |
| eDNA1706 | Hardluck-A | Piru River | 6/14/2021 | 34.68269 | -118.84377 | N | N |
| eDNA1707 | Hardluck-B | Piru River | 6/14/2021 | 34.68269 | -118.84377 | N | N |
| eDNA1708 | Hardluck-C | Piru River | 6/14/2021 | 34.68269 | -118.84377 | N | N |
| eDNA1709 | Sespe-Beaver-A | Sespe Creek | 6/14/2021 | 34.55422 | -119.24001 | N | N |
| eDNA1710 | Sespe-Beaver-B | Sespe Creek | 6/14/2021 | 34.55422 | -119.24001 | N | N |
| eDNA1711 | Sespe-Beaver-C | Sespe Creek | 6/14/2021 | 34.55422 | -119.24001 | N | N |
| eDNA1712 | Sespe-lion 1-A | Sespe Creek | 6/15/2021 | 34.56174 | -119.16302 | N | N |
| eDNA1713 | Sespe-lion 1-B | Sespe Creek | 6/15/2021 | 34.56174 | -119.16302 | N | N |
| eDNA1714 | Sespe-lion 1-C | Sespe Creek | 6/15/2021 | 34.56174 | -119.16302 | N | N |
| eDNA1633 | Sespe-lion 2-A | Sespe Creek | 6/15/2021 | 34.55958 | -119.16114 | N | N |
| eDNA1634 | Sespe-lion 2-B | Sespe Creek | 6/15/2021 | 34.55958 | -119.16114 | N | N |
| eDNA1635 | Sespe-lion 2-C | Sespe Creek | 6/15/2021 | 34.55958 | -119.16114 | N | N |
| eDNA1639 | Lower Piru 1-A | Piru River | 6/15/2021 | 34.52619 | -118.75694 | N | N |
| eDNA1640 | Lower Piru 1-B | Piru River | 6/15/2021 | 34.52619 | -118.75694 | N | N |
| eDNA1641 | Lower Piru 1-C | Piru River | 6/15/2021 | 34.52619 | -118.75694 | N | N |
| eDNA1642 | Lower Piru 2-A | Piru River | 6/16/2021 | 34.53548 | -118.75904 | N | N |
| eDNA1643 | Lower Piru 2-B | Piru River | 6/16/2021 | 34.53548 | -118.75904 | N | N |
| eDNA1645 | Lower Piru 2-C | Piru River | 6/16/2021 | 34.53548 | -118.75904 | N | N |
| eDNA1680 | Hardluck02-A | Piru River | 7/14/2021 | 34.68355 | -118.84421 | N | N |
| eDNA1681 | Hardluck02-B | Piru River | 7/14/2021 | 34.68355 | -118.84421 | N | N |
| eDNA1693 | Hardluck02-C | Piru River | 7/14/2021 | 34.68355 | -118.84421 | N | N |
| eDNA1716 | 8C-a | Santa Margarita River | 5/10/2022 | 33.31084 | -117.34658 | Y | N |
| eDNA1717 | 8C-b | Santa Margarita River | 5/10/2022 | 33.31084 | -117.34658 | Y | N |
| eDNA1718 | 8C-c | Santa Margarita River | 5/10/2022 | 33.31084 | -117.34658 | Y | N |
| eDNA1719 | 9D-a | Santa Margarita River | 5/10/2022 | 33.32325 | -117.33677 | Y | N |
| eDNA1720 | 9D-b | Santa Margarita River | 5/10/2022 | 33.32325 | -117.33677 | Y | N |
| eDNA1721 | 9D-c | Santa Margarita River | 5/10/2022 | 33.32325 | -117.33677 | Y | Y |
| eDNA1722 | 9E-a | Santa Margarita River | 5/10/2022 | 33.32495 | -117.33669 | Y | N |
| eDNA1723 | 9E-b | Santa Margarita River | 5/10/2022 | 33.32495 | -117.33669 | Y | Y |
| eDNA1724 | 9E-c | Santa Margarita River | 5/10/2022 | 33.32495 | -117.33669 | Y | Y |
| eDNA1725 | 10B-a | Santa Margarita River | 5/10/2022 | 33.33096 | -117.33334 | Y | Y |
| eDNA1726 | 10B-b | Santa Margarita River | 5/10/2022 | 33.33096 | -117.33334 | Y | Y |
| eDNA1727 | 10B-c | Santa Margarita River | 5/11/2022 | 33.33096 | -117.33334 | Y | Y |
| eDNA1728 | 10D-a | Santa Margarita River | 5/10/2022 | 33.33809 | -117.33151 | Y | N |
| eDNA1729 | 10D-b | Santa Margarita River | 5/10/2022 | 33.33809 | -117.33151 | Y | N |
| eDNA1730 | 10D-c | Santa Margarita River | 5/10/2022 | 33.33809 | -117.33151 | Y | N |
| eDNA1732 | 11F-a | Santa Margarita River | 5/11/2022 | 33.35205 | -117.32756 | Y | Y |
| eDNA1733 | 11F-b | Santa Margarita River | 5/11/2022 | 33.35205 | -117.32756 | Y | Y |
| eDNA1734 | 11F-c | Santa Margarita River | 5/11/2022 | 33.35205 | -117.32756 | Y | Y |
| eDNA1735 | 12A-a | Santa Margarita River | 5/11/2022 | 33.35277 | -117.32672 | Y | N |
| eDNA1736 | 12A-b | Santa Margarita River | 5/11/2022 | 33.35277 | -117.32672 | Y | Y |
| eDNA1737 | 12A-c | Santa Margarita River | 5/11/2022 | 33.35277 | -117.32672 | Y | Y |
| eDNA1738 | 12C-a | Santa Margarita River | 5/11/2022 | 33.35555 | -117.32583 | Y | N |
| eDNA1739 | 12C-b | Santa Margarita River | 5/11/2022 | 33.35555 | -117.32583 | Y | N |
| eDNA1740 | 12C-c | Santa Margarita River | 5/11/2022 | 33.35555 | -117.32583 | Y | N |

**Table S7.** eDNA samples collected for field validation of our species-specific qPCR assay for *Spea hammondii.*

| **Sample name** | **Site name** | **Location** | **Sample collection date** | **Latitude** | **Longitude** | ***S. hammondii* visually observed** | ***S. hammondii* detected with qPCR** |
| --- | --- | --- | --- | --- | --- | --- | --- |
| eDNA1091 | FHL_LK_49_SITE1 | Fort Hunter Ligget | 3/29/2019 | 35.94575 | -121.16048 | Y | Y |
| eDNA1170 | FHL_LK_49_SITE1 | Fort Hunter Ligget | 3/29/2019 | 35.94575 | -121.16048 | Y | Y |
| eDNA1175 | FHL_LK_49_SITE1 | Fort Hunter Ligget | 3/29/2019 | 35.94575 | -121.16048 | Y | Y |
| eDNA1083 | FHL_LK_49_SITE2 | Fort Hunter Ligget | 3/29/2019 | 35.94712 | -121.15973 | Y | Y |
| eDNA1171 | FHL_LK_49_SITE2 | Fort Hunter Ligget | 3/29/2019 | 35.94712 | -121.15973 | Y | Y |
| eDNA1180 | FHL_LK_49_SITE2 | Fort Hunter Ligget | 3/29/2019 | 35.94712 | -121.15973 | Y | Y |
| eDNA1136 | FHL_LK_49_SITE3 | Fort Hunter Ligget | 3/29/2019 | 35.9481 | -121.16034 | Y | Y |
| eDNA1141 | FHL_LK_49_SITE3 | Fort Hunter Ligget | 3/29/2019 | 35.9481 | -121.16034 | Y | Y |
| eDNA1146 | FHL_LK_49_SITE3 | Fort Hunter Ligget | 3/29/2019 | 35.9481 | -121.16034 | Y | Y |
| eDNA936 | FHL_LK_51_SITE1 | Fort Hunter Ligget | 3/29/2019 | 35.95024 | -121.16005 | N | N |
| eDNA947 | FHL_LK_51_SITE1 | Fort Hunter Ligget | 3/29/2019 | 35.95024 | -121.16005 | N | N |
| eDNA958 | FHL_LK_51_SITE1 | Fort Hunter Ligget | 3/29/2019 | 35.95024 | -121.16005 | N | N |
| eDNA937 | FHL_LK_51_SITE2 | Fort Hunter Ligget | 3/29/2019 | 35.95135 | -121.15971 | N | N |
| eDNA948 | FHL_LK_51_SITE2 | Fort Hunter Ligget | 3/29/2019 | 35.95135 | -121.15971 | N | N |
| eDNA959 | FHL_LK_51_SITE2 | Fort Hunter Ligget | 3/29/2019 | 35.95135 | -121.15971 | N | N |
| eDNA998 | FHL_LK_78_SITE1 | Fort Hunter Ligget | 4/7/2019 | 36.07509 | -121.26655 | N | Y |
| eDNA1094 | FHL_LK_78_SITE1 | Fort Hunter Ligget | 4/7/2019 | 36.07509 | -121.26655 | N | Y |
| eDNA1103 | FHL_LK_78_SITE1 | Fort Hunter Ligget | 4/7/2019 | 36.07509 | -121.26655 | N | Y |
| eDNA972 | FHL_LK_78_SITE2 | Fort Hunter Ligget | 4/7/2019 | 36.07488 | -121.265605 | N | Y |
| eDNA999 | FHL_LK_78_SITE2 | Fort Hunter Ligget | 4/7/2019 | 36.07488 | -121.265605 | N | Y |
| eDNA1095 | FHL_LK_78_SITE2 | Fort Hunter Ligget | 4/7/2019 | 36.07488 | -121.265605 | N | Y |
| eDNA1116 | FHL_LK_78_SITE3 | Fort Hunter Ligget | 4/7/2019 | 36.07482 | -121.26525 | N | Y |
| eDNA1104 | FHL_LK_78_SITE3 | Fort Hunter Ligget | 4/7/2019 | 36.07482 | -121.26525 | N | N |
| eDNA1657 | Homestead-A | MCAS Miramar | 3/22/2021 | 32.88544 | -117.0775 | N | N |
| eDNA1658 | Homestead-B | MCAS Miramar | 3/22/2021 | 32.88544 | -117.0775 | N | N |
| eDNA1664 | Homestead-C | MCAS Miramar | 3/22/2021 | 32.88544 | -117.0775 | N | N |
| eDNA1671 | RC01-A | Rancho Cebada | 4/21/2021 | 34.69811 | -120.36649 | N | N |
| eDNA1672 | RC01-B | Rancho Cebada | 4/21/2021 | 34.69811 | -120.36649 | N | N |
| eDNA1673 | RC01-C | Rancho Cebada | 4/21/2021 | 34.69811 | -120.36649 | N | N |
| eDNA1659 | RC02-A | Rancho Cebada | 4/21/2021 | 34.70534 | -120.37629 | N | N |
| eDNA1667 | RC02-B | Rancho Cebada | 4/21/2021 | 34.70534 | -120.37629 | N | N |
| eDNA1683 | RC02-C | Rancho Cebada | 4/21/2021 | 34.70534 | -120.37629 | N | N |
| eDNA1674 | RC03-A | Rancho Cebada | 4/21/2021 | 34.70616 | -120.367 | N | N |
| eDNA1675 | RC03-B | Rancho Cebada | 4/21/2021 | 34.70616 | -120.367 | N | N |
| eDNA1676 | RC03-C | Rancho Cebada | 4/21/2021 | 34.70616 | -120.367 | N | N |
| eDNA1677 | RC04-A | Rancho Cebada | 4/21/2021 | 34.70665 | -120.34728 | Y | N |
| eDNA1678 | RC04-B | Rancho Cebada | 4/21/2021 | 34.70665 | -120.34728 | Y | N |
| eDNA1679 | RC04-C | Rancho Cebada | 4/21/2021 | 34.70665 | -120.34728 | Y | N |
| eDNA1660 | TG_5014-A | MCBCP | 3/23/2021 | 33.20552 | -117.46005 | Y | Y |
| eDNA1665 | TG_5014-B | MCBCP | 3/23/2021 | 33.20552 | -117.46005 | Y | N |
| eDNA1666 | TG_5014-C | MCBCP | 3/23/2021 | 33.20552 | -117.46005 | Y | Y |
| eDNA1689 | TG_5040-A | MCBCP | 3/23/2021 | 33.30495 | -117.45932 | Y | Y |
| eDNA1690 | TG_5040-B | MCBCP | 3/23/2021 | 33.30495 | -117.45932 | Y | Y |
| eDNA1691 | TG_5040-C | MCBCP | 3/23/2021 | 33.30495 | -117.45932 | Y | Y |
| eDNA1662 | TG_75009-A | MCBCP | 3/23/2021 | 33.3061 | -117.46053 | Y | Y |
| eDNA1669 | TG_75009-B | MCBCP | 3/23/2021 | 33.3061 | -117.46053 | Y | Y |
| eDNA1685 | TG_75009-C | MCBCP | 3/23/2021 | 33.3061 | -117.46053 | Y | Y |
| eDNA1686 | TG_75016-A | MCBCP | 3/23/2021 | 33.30502 | -117.45992 | Y | Y |
| eDNA1687 | TG_75016-B | MCBCP | 3/23/2021 | 33.30502 | -117.45992 | Y | Y |
| eDNA1688 | TG_75016-C | MCBCP | 3/23/2021 | 33.30502 | -117.45992 | Y | Y |
| eDNA1661 | Willow01-A | Willow Ranch | 4/21/2021 | 34.69582 | -120.35518 | Y | Y |
| eDNA1668 | Willow01-B | Willow Ranch | 4/21/2021 | 34.69582 | -120.35518 | Y | Y |
| eDNA1684 | Willow01-C | Willow Ranch | 4/21/2021 | 34.69582 | -120.35518 | Y | Y |

**Table S8.** GenBank sequences identified by Primer-BLAST with the potential to cross-amplify with the qPCR assay that we designed for *Spea hammondii.* None of the sequences belong to species that are sympatric with *Spea hammondii.*

| **Genbank accesion #** | **Species** | **Description** |
| --- | --- | --- |
| EF508476.1 | *Acanthopsoides hapalias* | Asian freshwater fish species |
| HM536795.1 | *Acrossocheilus monticola* | Asian freshwater fish species |
| KC696534.1 | *Acrossocheilus yunnanensis* | Asian freshwater fish species |
| KF113879.1 | *Acrossocheilus barbodon* | Asian freshwater fish species |
| KJ994678.1 | *Acrossocheilus iridescens* | Asian freshwater fish species |
| FJ752340.1 | *Ameerega smaragdina* | South American anuran species |
| JQ012377.1 | *Anchoa parva* | Marine fish species |
| JQ012387.1 | *Anchoa filifera* | Marine fish species |
| JQ012327.1 | *Anchoviella cf.* | Marine fish species |
| AB279333.1 | *Anguilla australis* | Australian freshwater eel species |
| MH800477.1 | *Anguilla mossambica* | African freshwater eel species |
| GU932709.1 | *Artibeus obscurus* | South American bat species |
| AF334087.1 | *Barbus graellsii* | European freshwater fish species |
| AY887784.1 | *Botia striata* | Asian freshwater fish species |
| KY065276.1 | *Capoeta capoeta* | Asian freshwater fish species |
| AF187018.1 | *Carollia brevicauda* | South American bat species |
| AF187020.1 | *Carollia castanea* | South American bat species |
| KC631284.1 | *Cirrhinus molitorella* | Asian freshwater fish species |
| AF475153.1 | *Clarias gariepinus* | African freshwater fish species |
| EF422294.1 | *Coilia mystus* | Marine fish species |
| AP009581.1 | *Conocara murrayi* | Marine fish species |
| AY953001.1 | *Coreius guichenoti* | Asian freshwater fish species |
| AP012148.1 | *Crossocheilus latius* | Asian freshwater fish species |
| AP013321.1 | *Crossocheilus langei* | Asian freshwater fish species |
| JX074225.1 | *Crossocheilus oblongus* | Asian freshwater fish species |
| JX083158.1 | *Crossocheilus diplochilus* | Asian freshwater fish species |
| DQ422087.1 | *Galeorhinus galeus* | Marine fish species |
| MH500684.1 | *Garra cambodgiensis* | Asian freshwater fish species |
| EU552578.1 | *Gilchristella aestuaria* | African freshwater fish species |
| AF416891.1 | *Glyptosternon maculatum* | Asian freshwater fish species |
| MT606734.1 | *Gymnocranius frenatus* | Marine fish species |
| AP012012.1 | *Hara jerdoni* | Asian freshwater fish species |
| HQ257283.1 | *Hemibagrus filamentus* | Asian freshwater fish species |
| KJ573466.1 | *Hemibagrus nemurus* | Asian freshwater fish species |
| KJ624624.1 | *Hemibagrus wyckioides* | Asian freshwater fish species |
| AP011386.1 | *Henicorhynchus lineatus* | Asian freshwater fish species |
| AP009582.1 | *Herwigia kreffti* | Marine fish species |
| KJ158097.1 | *Ilisha amazonica* | South American freshwater fish species |
| MG958199.1 | *Ilisha melastoma* | Marine fish species |
| KY132250.1 | *Kaloula conjuncta* | Asian anuran species |
| EF508526.1 | *Kottelatlimia pristes* | Asian freshwater fish species |
| GQ853086.1 | *Labeo fimbriatus* | Asian freshwater fish species |
| MH026027.1 | *Labeo lankae* | Asian freshwater fish species |
| MH026035.1 | *Labeo porcellus* | Asian freshwater fish species |
| KT698047.1 | *Lamiopsis tephrodes* | Marine fish species |
| KT698048.1 | *Lamiopsis temminckii* | Marine fish species |
| AB080154.1 | *Lefua echigonia* | Asian freshwater fish species |
| EU036445.1 | *Lepidorhombus whiffiagonis* | Marine fish species |
| FJ155494.1 | *Lonchorhina aurita* | South American bat species |
| EU932868.1 | *Macropodus hongkongensis* | Asian freshwater fish species |
| AB239569.1 | *Mantella bernhardi* | African anuran species |
| AY157035.1 | *Mesophylla macconnelli* | South American Bat species |
| HM142578.1 | *Onychostoma lepturum* | Asian freshwater fish species |
| HQ235764.1 | *Onychostoma rara* | Asian freshwater fish species |
| JX074246.1 | *Onychostoma ovale* | Asian freshwater fish species |
| AP004431.1 | *Ostichthys japonicus* | Marine fish species |
| MT850132.1 | *Parabotia kiangsiensis* | Asian freshwater fish species |
| AP009619.1 | *Pellona flavipinnis* | South American freshwater fish species |
| EU552554.1 | *Pellona castelnaeana* | South American freshwater fish species |
| AF201604.1 | *Petrocephalus simus* | African freshwater fish species |
| EU770156.1 | *Petrocephalus odzalaensis* | African freshwater fish species |
| EU770183.1 | *Petrocephalus christyi* | African freshwater fish species |
| EU770191.1 | *Petrocephalus balayi* | African freshwater fish species |
| GU982923.1 | *Petrocephalus stuhlmanni* | African freshwater fish species |
| KJ409658.1 | *Petrocephalus wesselsi* | African freshwater fish species |
| KJ409663.1 | *Petrocephalus catostoma* | African freshwater fish species |
| GQ244477.1 | *Pipa pipa* | South American anuran species |
| MT157615.1 | *Prolixicheilus longisulcus* | Asian freshwater fish species |
| AY912410.1 | *Pseudobagrus pratti* | Asian freshwater fish species |
| EU241453.1 | *Puntius denisonii* | Asian freshwater fish species |
| KF574537.1 | *Rasbora daniconius* | Asian freshwater fish species |
| GQ406329.1 | *Rectoris posehensis* | Asian freshwater fish species |
| KF410777.1 | *Rhodeus aff.* | Asian freshwater fish species |
| AF472583.1 | *Sardinella maderensis* | Marine fish species |
| FJ463052.1 | *Scutiger boulengeri* | Asain anuran species |
| FJ945491.1 | *Scutiger tuberculatus* | Asain anuran species |
| FJ945495.1 | *Scutiger mammatus* | Asain anuran species |
| KC140561.1 | *Scutiger ningshanensis* | Asain anuran species |
| EF191043.1 | *Spea bombifrons* | North American anuran species, not sympatric with *A. californicus* or *S. hammondii* |
| AF435136.1 | *Sturnira lilium* | South American bat species |
| AF435170.1 | *Sturnira luisi* | South American bat species |
| AF435175.1 | *Sturnira ludovici* | South American bat species |
| AF435177.1 | *Sturnira magna* | South American bat species |
| AF435184.1 | *Sturnira tildae* | South American bat species |
| AF435206.1 | *Sturnira oporaphilum* | South American bat species |
| AF435212.1 | *Sturnira mordax* | South American bat species |
| AF435240.1 | *Sturnira erythromos* | South American bat species |
| AF435246.1 | *Sturnira bogotensis* | South American bat species |
| AF435252.1 | *Sturnira aratathomasi* | South American bat species |
| KC753793.1 | *Sturnira hondurensis* | South American bat species |
| KY366229.1 | *Sturnira adrianae* | South American bat species |
| KP670368.1 | *Tachysurus adiposalis* | Asian freshwater fish species |
| MN585245.1 | *Tonatia saurophila* | South American bat species |
| AF435173.1 | *Vampyressa brocki* | South American bat species |
| NC_044879.1 | *Xenopus itombwensis* | African anuran species |

**Appendix I**. Modified protocol for eDNA filter extraction.

For extraction of eDNA filters, we used the Qiagen DNeasy blood and tissue kit, with the following modifications to the standard protocol:

Extractions were carried out in a room dedicated to only eDNA extractions, under a laminar flow PCR hood with UV sterilization. For samples where 5 L of water was filtered, we extracted one-half of each filter. For filters where less than 5 L of water was filtered, we extracted the entire filter.

Filters were first cut in half, and each half filter was cut into 5 pieces using sterilized forceps and scissors, inside a sterile petri dish. The cut filter pieces were then placed in 1.5 ml microcentrifuge tubes already containing 360 µL of buffer ATL and 40 µL proteinase K. Tubes were then incubated for 18-36 hours on a shaking incubator at 56° C.

Following incubation, we added 400 µL of buffer AL, vortexed for 30 seconds, and centrifuged for two minutes at maximum speed to push liquid down into the filter. Tubes were then incubated for 10 minutes at 70° on a shaking incubator. We then added 400 µL 100% ethanol, vortexed for 30 seconds, and centrifuged for two minutes at maximum speed to push liquid down into the filter. We then moved 700 µL of supernatant from each sample into Qiagen DNeasy Mini spin columns, centrifuged them for one minute at maximum speed, and placed the filter columns into new collection tubes. Next, we moved all the filter material from the initial incubation tubes into Qiagen QIAshredder spin columns with sterilized forceps, and centrifuged the QIAshredder spin columns for two minutes at maximum speed. We then moved the supernatant from the QIAshredder columns (liquid “sucked” out of the filters) to their respective Qiagen DNeasy Mini spin columns, centrifuged the spin columns for one minute at maximum speed, and placed them into new collection tubes.

For ‘whole filter’ extractions, these steps were then repeated with the second half of the filter, using the same Qiagen DNeasy Mini spin column as the first half of the filter.

We then proceeded with the wash and elution steps outlined in the standard Qiagen DNeasy protocol, with a final elution volume of 100 µL.
